# A Digital Twin to Optimize Treatment Efficacy of Targeted Alpha-particle Therapies by Antibody-Radioconjugate Cocktails Against Solid Tumors

**DOI:** 10.64898/2025.12.19.695379

**Authors:** Michail Kavousanakis, Rachel Macher, Yannis Kevrekidis, Stavroula Sofou

## Abstract

Advanced solid tumors are incurable. Antibody-delivered targeted alpha-particle (α-particle) radionuclide therapies (TAT) comprise a tumor-agnostic treatment type, due to the unparalleled killing efficacy of, and irradiation precision (4-5 cell lengths) by, α-particles, as well as the selectivity in tumor cell targeting by antibody technologies. However, cells not being directly hit by α-particles will likely not be killed.

**METHODS:** To address the limited solid tumor penetration by highly specific and strongly binding antibody-radioconjugates, an experimentally informed digital twin, based on transport (diffusion/advection) first principles, was developed to describe an approach where a fraction of the administered (radio)activity is delivered by a separate type of a model α-particle antibody-radioconjugate of low(er)/no affinity for the same marker. The latter was chosen because it can irradiate cells residing in the deep regions of solid tumors away from vasculature.

**RESULTS:** The digital twin that was trained and validated on spheroids that were employed as surrogates of the avascular tumor regions, demonstrated that the investigated cocktails of antibody-radioconjugates with controlled affinities exhibited better inhibition of spheroid growth compared to the extent of growth inhibition by the high-affinity antibody-radioconjugate alone, for the same total (incubated) activity concentrations; this prediction was independent of spheroid size and/or level of expression of the targeted markers.

**CONCLUSION:** The findings of this study suggest that antibody-delivered TAT (that is already in the clinic) can be augmented by delivering a fraction of the same total activity by low(er) affinity antibody-radioconjugates. This combination of separate antibody-radioconjugates with variable affinities (for the same targeted marker) is a promising approach to possibly even more delay recurrence and further prolong survival of patients with advanced solid tumors.

## Introduction

Advanced solid tumors are difficult to treat and are commonly incurable. In 2025, the American Cancer Society estimated that over 618,000 patients will die from cancer in the US, and a conservative minimum of 50% of these cases will involve incurable solid tumors ^1^. Targeted alpha-particle (α-particle) radionuclide therapeutics (TAT) – that are being investigated as alternative treatments - are presently experiencing a renaissance: from one clinical trial in 2016 to 20 active clinical trials in 2024 in the US alone, with 70% involving tumor-targeting antibody-radioconjugates being evaluated against tumors of different origins. The enthusiasm for TAT, which is arguably a tumor-agnostic treatment, is due to the unparalleled killing efficacy of, and irradiation precision by, α-particles, as well as to the selectivity in tumor targeting by antibody technologies ^2^. α-particles travel only 4-5 cell lengths in tissue and act like bullets, causing complex double-strand DNA breaks as they traverse the cell nucleus. The inability to repair this DNA damage ^3^ is the reason that α-particles are impervious to resistance ^4–6^, independent of cell origin and/or resistance to other agents ^4^.

However, cells not being directly hit by α-particles will likely not be killed. Antibody-radioconjugates are successful in selectively delivering high doses to tumors, but the more strongly an antibody binds on cancer cells, the less deep it penetrates into solid tumors ^7,8^, resulting in fewer cancer cells effectively being irradiated and killed deeper in the tumor. This is the main reason why patients treated with TAT often ultimately relapse ^9^: cells in deep tumor regions far from vasculature often do not receive lethal doses of antibody-delivered agents injected in the blood.

To address the limited solid tumor penetration by highly specific and strongly binding antibody-radioconjugates, we employed an additional separate type of a model α-particle antibody-radioconjugate of low(er)/no affinity for the same marker. The low(er)/no affinity antibody-radioconjugates were chosen because they can irradiate cells residing in the deep regions of solid tumors away from vasculature, since their diffusion-limited transport is not obstructed as much by their binding to cancer cell markers, as that of the high-affinity antibody-radioconjugates’.

A digital twin was developed to describe the cocktails of these two types of high and low(er)/no affinity antibody-radioconjugates with the goal of identifying the optimal (radio)activity split ratios between the two antibody types; the goal was to minimize the cell survival fraction in spheroids of different sizes that were employed as surrogates of the avascular tumor regions.

## Results

We discuss three types of simulated therapeutic treatment scenarios, schematically illustrated in Fig.1. The first two scenarios involve treatments based on a single type of carrier: either Trastuzumab alone or solely Rituximab (top and middle panels in Fig.1, respectively). In the third scenario, we simulate the combined action of Trastuzumab and Rituximab using a “carrier-cocktail”. In all scenarios, the spheroids are initially immersed in a surrounding medium with a constant concentration of carriers for a time interval 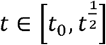, corresponding to the incubation stage. Afterwards, the spheroids are immersed in a carrier-free medium, marking the clearance stage.

**Fig. 1.**
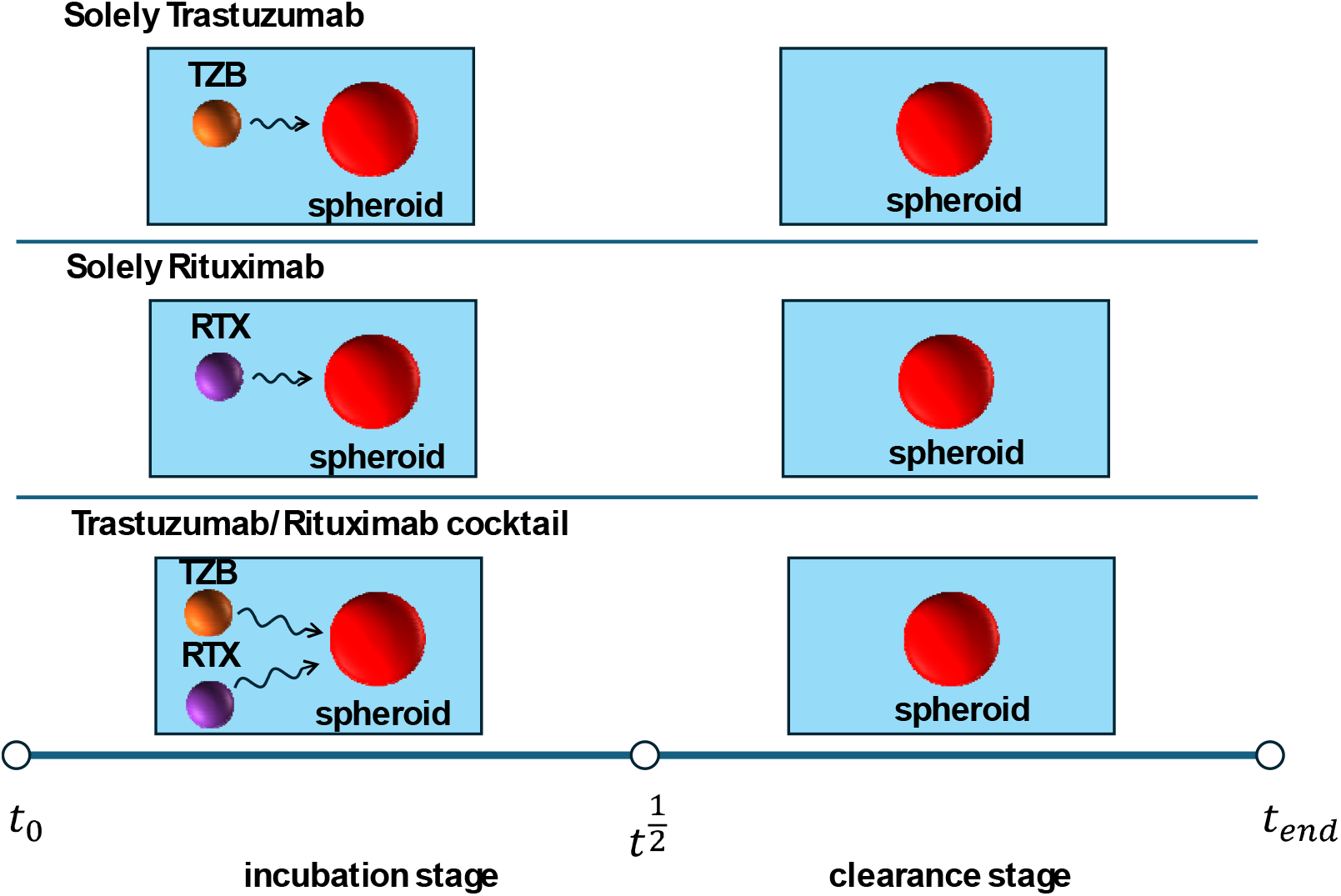
Schematic representation of the administration schedules used in simulations.

### Single-carrier simulations

We first present results from single-carrier simulations. Tumor spheroids are incubated for 22 hrs in a solution containing either Trastuzumab-radioconjugate or Rituximab-radioconjugate at constant concentration, and killing efficacy is evaluated 24 hrs after the end of the incubation period.

All simulations assume the same external carrier concentrations and a fixed isotope-to-antibody ratio, *a*_*r*_. Figure 2 (A)&(B) show the spatial distribution of [^225^Ac] at the end of incubation and after 24 hrs of clearance, using Trastuzumab (TZB) or Rituximab (RTX). The isotope concentration in the surrounding solution is set to 0.0019 nM (or equivalently 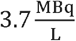).

**Fig. 2.**
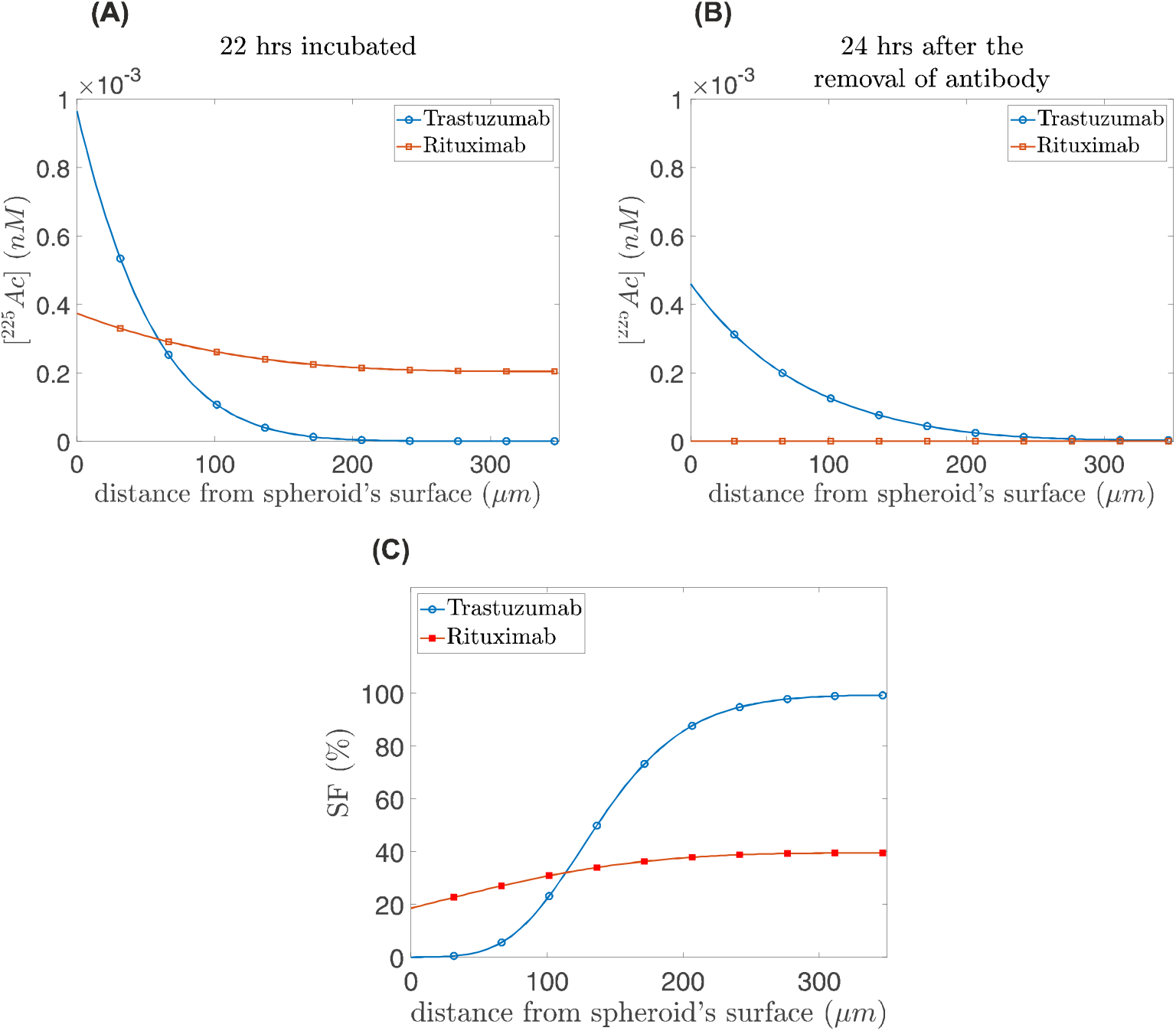
Single carrier simulations. Isotope concentration profiles in a spheroid of 350 μm radius: **(A)** at the end of the incubation period (22 hrs) and **(B)** after a 24-hour clearance period. **(C)** Survival fraction of cancer cells after a 24-hr clearance period as a function of distance from spheroid’s surface. Results are shown for Trastuzumab-radioconjugates, and Rituximab-radioconjugates.

Due to its high affinity for cancer cell receptors, Trastuzumab delivers a higher isotope concentration to the spheroid periphery but exhibits limited penetration into deeper regions. In contrast, Rituximab penetrates more deeply into the spheroid core; however, due to its “non-stickiness” to cancer cells, the delivered isotope is rapidly cleared, resulting in negligible concentrations 24 hrs post-incubation.

Administration schedules used in simulations with Trastuzumab (high affinity antibody-radioconjugate) and Rituximab (no affinity antibody-radioconjugate) delivering the isotope cargo during the incubation stage 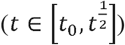. The incubation stage is followed by a clearance period in which the spheroids are immersed in a carrier-free surrounding medium. In all cases, the incubation period ends at 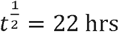 (clearance time of Trastuzumab and Rituximab).

Consequently, Trastuzumab-radioconjugates achieve strong cytotoxicity near the spheroid surface, whereas Rituximab-radioconjugates distribute the isotope more uniformly and target cells also in the interior. This trend is reflected in the spatial distribution of cancer cell survival fraction (see Fig.2(C)). Trastuzumab-radioconjugates yield near-complete killing at the spheroid periphery (*SF* ≈ 0) but rapidly loses efficacy toward the center. Rituximab-radioconjugates, while less effective at the surface, exhibit enhanced killing in deeper regions.

Survival fractions (*SF*) are computed using equation (4) with 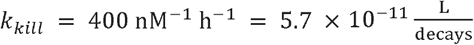.

### Cocktail-carrier simulations

Given that Rituximab penetrates more effectively into the spheroid core while Trastuzumab primarily targets the periphery, we hypothesize that combining the two carriers can enhance overall killing efficacy by producing a more uniform isotope distribution. To test this, we partition the total isotope concentration (0.0019 nM) such that a fraction *F* is delivered by Trastuzumab and 1-*F* by Rituximab.

### Effect of the isotope fraction delivered by Trastuzumab (high-affinity antibody-radioconjugate)

We examine the effect of varying *F* a spheroid of radius 350 µm. Figure 3(A) depicts the spatial distribution of ^225^Ac at the end of incubation for different values of *F*, alongside single-carrier cases. As the Rituximab-radioconjugate contribution increases, the isotope distribution becomes progressively more uniform, while the peak concentration near the spheroid surface decreases. Figure 3(B) illustrates the corresponding distributions 24 hrs after the completion of incubation. Although Rituximab initially delivers higher isotope concentrations in the spheroid interior, its non-binding nature leads to faster depletion compared to Trastuzumab-rich cocktails. These trends are reflected in the survival fraction (*SF*) profiles (Fig. 3(C)). High Trastuzumab-radioconjugate fractions result in near-complete eradication within ~50 µm of the spheroid surface but leave the core largely unaffected. Increasing the Rituximab-radioconjugate contribution reduces survival in the interior but compromises killing efficiency near the surface. The optimal balance is achieved for a cocktail containing 45% Trastuzumab-radioconjugates and 55% Rituximab-radioconjugates, which minimizes the average survival fraction to 21.5% (Fig.3(D)).

**Fig. 3.**
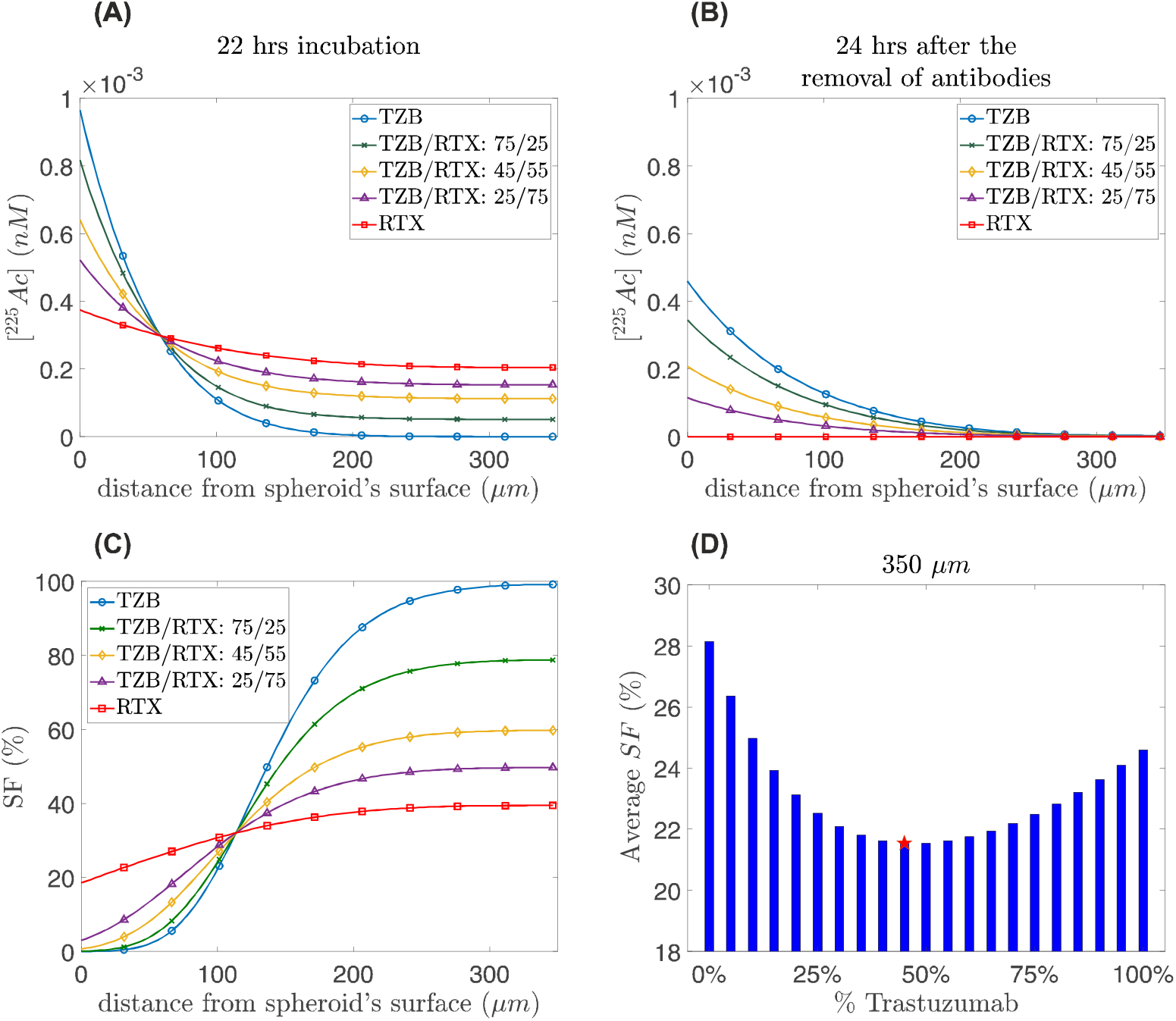
Carrier-cocktail simulations for a 350 μm radius spheroid. Distribution of isotope concentration throughout the spheroid at **(A)** the end of incubation (22 hrs), and **(B)** 24 hrs post-incubation. For comparison, we illustrate also distributions of isotopes carried by solely Trastuzumab and Rituximab. **(C)** Distribution of *SF* 24 hrs post-incubation for different cocktail treatments. **(D)** Average value of survival fraction (*SF*) various fraction values, *F*, of the isotope delivered by Trastuzumab in combination with Rituximab. The red star indicates the optimal TZB/RTX combination with the lowest average *SF* (21.5%) achieved using a 45% TZB / 55% RTX cocktail.

### Effect of receptor expression level

We next investigate how receptor expression, quantified by the total receptor concentration, *R*_*T*_ influences therapeutic efficacy. Figure 4 shows the average survival fraction for different *F* and *R*_*T*_ values in a 350 μm spheroid. As receptor expression decreases, the optimal killing efficacy shifts toward higher Trastuzumab-radioconjugate fractions. For *R*_*T*_ =200 nM, the lowest survival fraction(≈ 12.5%) is achieved with 100% Trastuzumab delivery, reflecting enhanced penetration due to reduced binding-site saturation. Conversely, for high receptor expression (*R*_*T*_ = 1000 nM), optimal efficacy is obtained with lower Trastuzumab contribution; in this case, a cocktail containing 25% Trastuzumab yields the lowest survival fraction (≈ 24.4%).

**Fig. 4.**
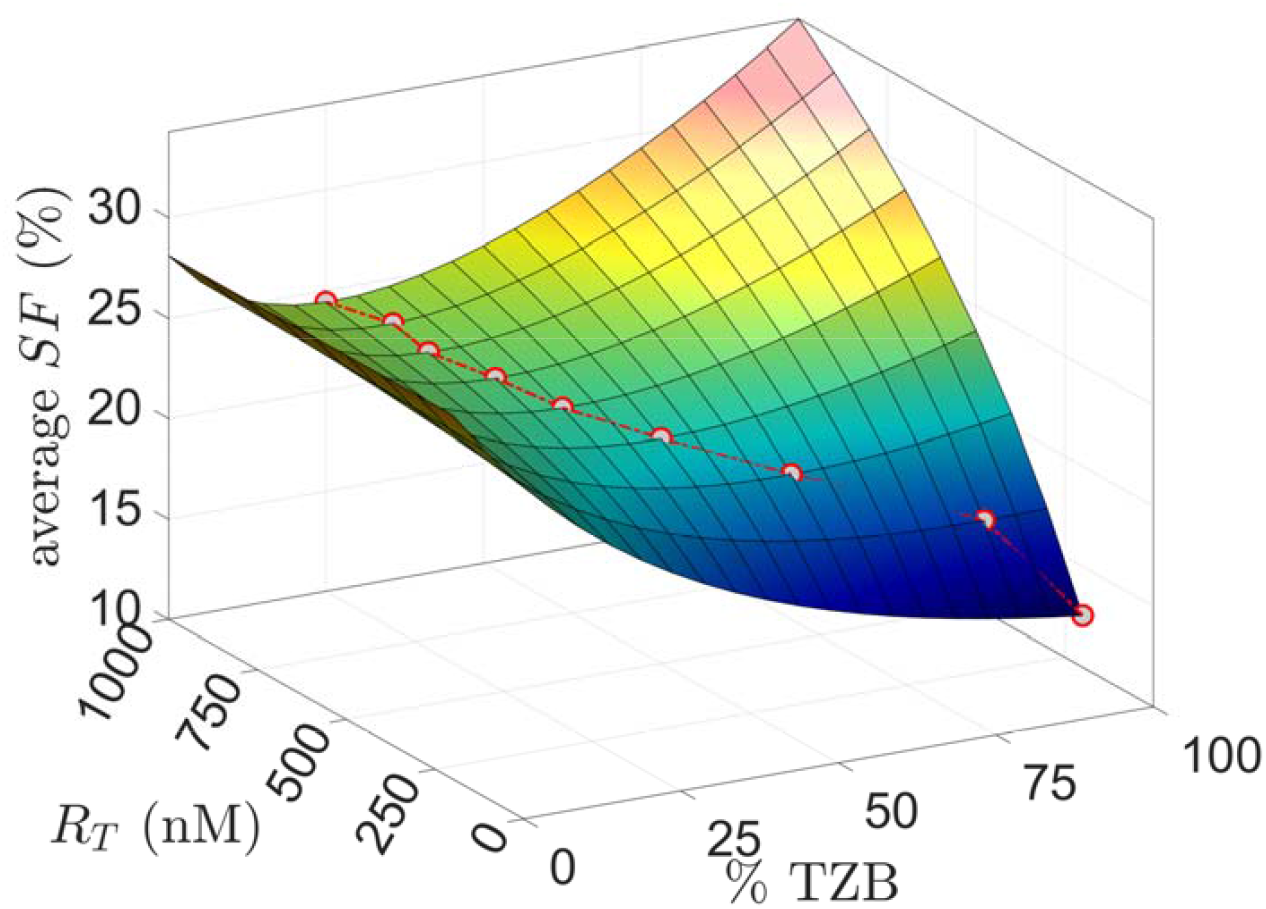
Effect of receptor expression level. Average survival fraction () for various fraction values of the isotope delivered by TZB and concentration of total receptors. The red-lined circles denote the optimum for each level of expression. Simulations are performed for a spheroid of 350 μm radius.

### Effect of spheroid size

Finally, we examine the impact of spheroid size - employed as the tumors’ avascular regions - on optimal cocktail composition for (see Fig.5). Smaller spheroids exhibit lower cell survival fractions and favor higher Trastuzumab-radioconjugate fractions, with near-complete eradication (=4.4%) achieved in 200 μm spheroids using Trastuzumab-radioconjugates alone. In contrast, larger spheroids show reduced overall cell death (greater cell survival fractions) and benefit from increased Rituximab-radioconjugate contribution, which improves isotope penetration into the spheroid core.

**Fig. 5.**
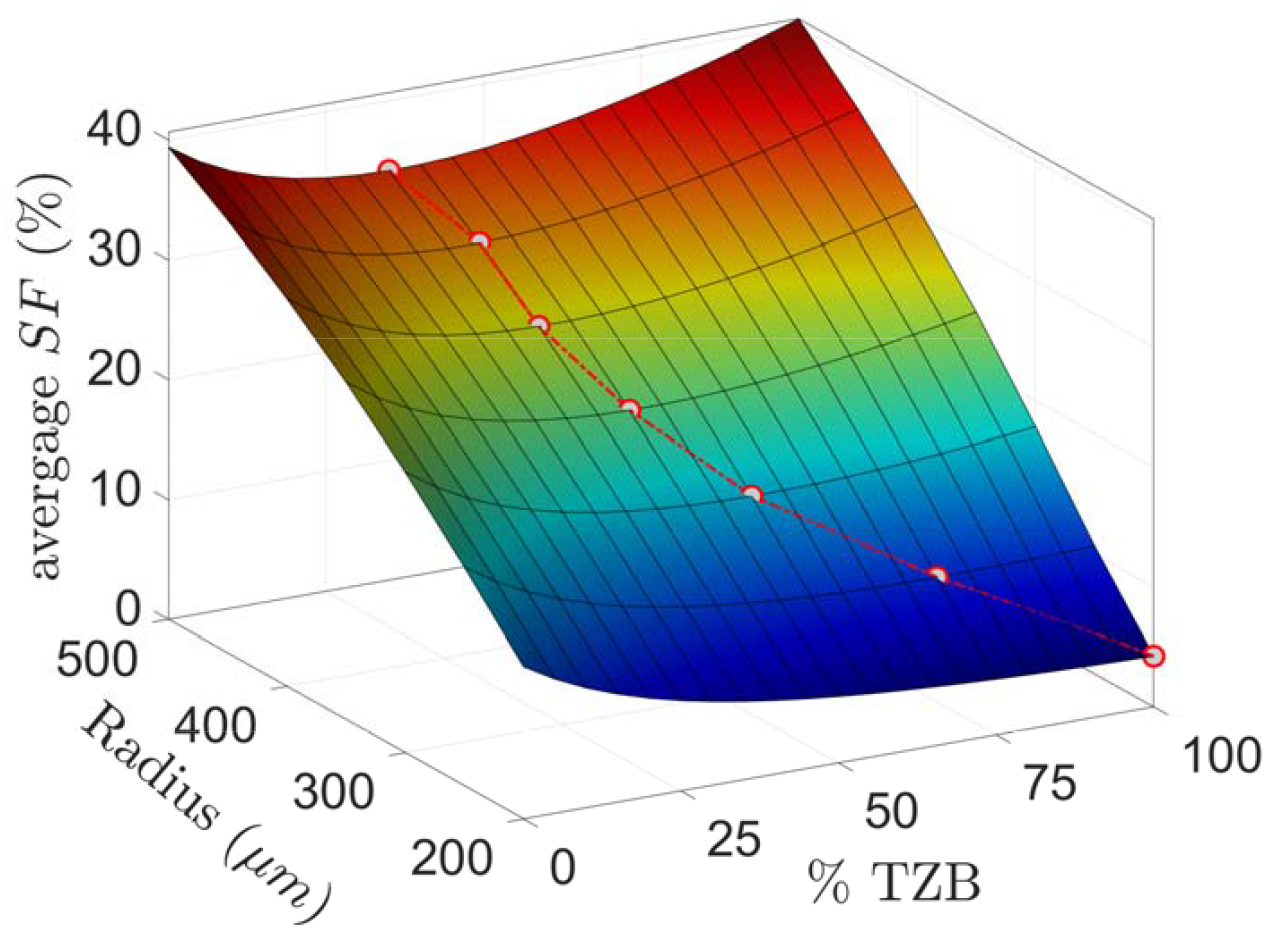
Effect of spheroid size. Average survival fraction () for various activity split ratios of the isotope delivered by TZB acros varying spheroid radii. Red-lined circles indicate the optimum for eachspheroid radius. Simulation are performed for a fixed receptor concentration of of receptors.

## Discussion

The experimentally informed digital twin demonstrated that the efficacy of TAT delivered by targeting antibody-radioconjugates can be improved by co-delivering a fraction of the injected activity by a second antibody-radioconjugate with low(er)/no affinity; the latter component essentially transports said fraction of α-emitters into the deeper regions of solid tumors, where high-affinity antibody-radioconjugates cannot reach. The investigated cocktails of antibody-radioconjugates with controlled affinities exhibited better inhibition of spheroid growth compared to the extent of tumor inhibition by the high-affinity antibody-radioconjugate alone, for the same total (incubated) activity concentrations and was independent of spheroid size and/or level of expression of the targeted markers.

Notably, the optimal activity split ratio shifted toward the low(er)/no affinity antibody-radioconjugates for greater spheroid sizes (corresponding to longer distances of tumor avascular regions) which can be explained by the larger cell population over longer distances in the deeper ends of the larger spheroids. Conversely, decreasing marker expression levels by cancer cells shifted the optimal activity split ratio toward the high-affinity antibody to enable delivery of lethal activities on the cells closer the spheroids edges; in these areas, essentially, only the high-affinity antibodies can effectively deliver substantial levels of activity, and with lowering binding sites a greater population of high-affinity antibody-radioconjugates was required to deliver lethal activity levels per cell.

Interestingly, in the latter case and within reasonable delivered activities, spheroids with lower expression of targeted markers were associated with better overall cell kill (lower survival fractions), mostly attributed to the deeper penetration even of the high-affinity antibody-radioconjugates, compared at the same activity concentrations during incubation of spheroids with the radioconjugates. It is possible, therefore, that by “passivating” a fraction of the highly expressed targeting markers before administering TAT enables deeper penetration even of the high-affinity radioconjugates since the available non-associated targeting markers would, at least transiently, be reduced.

In these studies, we have not interrogated the effect of the order of introducing each modality. It is possible that if we first incubate the spheroids with high-affinity antibody-radioconjugates and then follow with the low(er)/no affinity antibody-radioconjugates to observe even greater penetration of the latter within the interstitium, potentially increasing the fraction of killed cells ^10^.

We have previously reported a digital twin that described a systemically injected carrier cocktail approach to address the non-uniform α-particle irradiation patterns by antibody-radioconjugates within solid tumors ^11^. In our original report, the diffusion-driven delivery process was again separated from the targeting-driven delivery prosses enabling us with independent design variables to engineer a treatment for more uniform solid tumor irradiation by α-particles. The experimental approach comprised (a) as in the current study, cell-targeting antibody-radioconjugates that primarily irradiated the tumor regions closer to the vasculature, due to their diffusion-limited penetration; and (b) differently than the current study, liposomes selectively adhering to the tumor extracellular matrix (ECM), loaded with the same α-particle emitter; these liposomes were designed to sense the slightly-acidic tumor interstitium and release their highly diffusing radioactive cargo within the tumor interstitial space, where – as we demonstrated – exhibited significant penetration within the deeper avascular regions of solid tumors. That digital twin was experimentally informed with data acquired using multicellular spheroids. The predictions of that digital twin were validated in mice with subcutaneous xenografts, and demonstrated that carrier cocktails were more successful in inhibiting tumor growth than delivering the same total injected activity by the targeting antibody-radioconjugate alone, even at lower tumor absorbed doses (since the injected activity in liposomal form exhibited lower tumor uptake than the activity delivered by the targeting antibodies) ^7,12^.

The present study aims to, primarily, accelerate the potential clinical implementation/validation of carrier cocktails. Given that liposome-delivered α-particle therapy is not yet in the clinic, we investigated carrier cocktails comprising only antibody-radioconjugates with high and low affinities for the same target on cancer cells.

The longer circulation times of the (low(er)/no affinity) antibodies in the blood stream compared to those of liposomes, and the significantly smaller molecular size, contributing to greater diffusivities than those of liposomes, were hypothesized to be key properties that should allow the deep penetration of tumors. The findings of this study, suggest that combining separate antibody-radioconjugates with different affinities for solid tumor cancer cells, are a promising approach to augment antibody-delivered TAT that is already in the clinic and can possibly delay recurrence and prolong survival.

## Methods

We employ a reaction-diffusion transport model to describe the spatiotemporal evolution of specific (high-affinity) and non-specific (low-affinity) antibody-radioconjugates, within the volume of a tumor spheroid. The model comprises two stages: **(1)** An incubation stage, during which the spheroid is immersed in a solution with a constant concentration of antibody-radioconjugates. Carriers enter the spheroid through its outer surface and diffuse through the tumor interstitium. Once inside, specific antibody-radioconjugates may bind to receptors on the cancer cell surface, dissociate from them, or undergo internalization. Non-specific/non-binding antibody-radioconjugates enter the spheroid and are transported solely by diffusion. **(2)** A clearance stage, during which the external carrier concentration is set to zero and residual transport and reactions proceed within the spheroid.

The governing equations for the carrier concentration dynamics are:

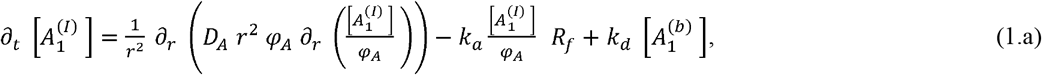

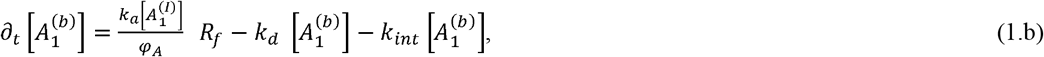

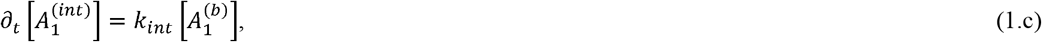

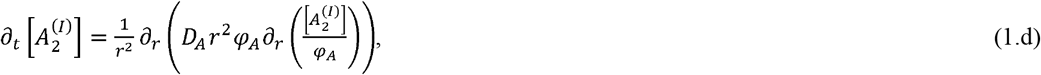

where 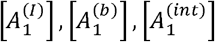 denote the concentrations of the specific antibody-radioconjugate in the interstitial space (denoted by ‘*I*’), bound to cell-surface receptors, and internalized within cells, respectively. 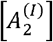 denotes the concentration of non-specific/non-binding antibody-radioconjugate, which exists only in the interstitial space. *D*_*A*_ is the effective diffusion coefficient of both carriers, and *φ*_*A*_ is the spheroid volume fraction accessible to antibodies *A*_1_,*A*_2_. The parameters *k*_*a*_, *k*_*d*_ and *k*_*int*_ represent the association (denoted by ‘*a*’), dissociation (denoted by ‘*d*’) and internalization (denoted by ‘*int*’) rate constants, respectively. *R*_*f*_ denotes the concentration of unoccupied/free cell-surface receptors available for binding with the specific antibody-radioconjugate, *A*_1_. We also consider the material balance for surface receptors, 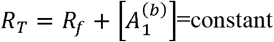, where *R*_*T*_ denotes the initial concentration of unbound receptors.

The carrier influx at the spheroid’s external surface is modeled using a Robin boundary condition:

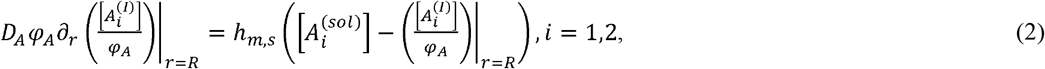

Where 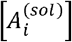 is the constant carrier concentration in the surrounding solution (*i*=1, for the specific antibody-radioconjugate, and *i* =2 for the non-specific antibody-radioconjugate); *h*_*m,s*_ is the mass transfer coefficient at the solution/spheroid interphase. This coefficient takes the values *h*_*m,inc*_ during incubation (when 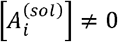) and *h*_*m,cl*_ during clearance (when 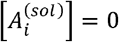).

In our simulations carrier 1 (specific antibody) corresponds to Trastuzumab (TZB, a HER2-targeting antibody), and carrier 2 to Rituximab (RTX, which does not recognize the HER2 receptor).

In prior work ^11^, transport and kinetic parameters, including apparent affinity constants and diffusivity for Trastuzumab and Rituximab, were estimated by fitting the reaction-diffusion model to experimental data. In the present study, we assume that Rituximab and Trastuzumab share identical transport properties (same diffusivities, mass transfer coefficients, and circulation times in the blood). All parameter values are summarized in Table 1.

**Table 1.** Transport and kinetic parameters for Trastuzumab and Rituximab^11,15^.

| Property | Value / Expression |
| --- | --- |
| Effective diffusion coefficient, $D_A$ | $6 \frac{\mu\text{m}^2}{\text{s}}$ |
| Porosity, $\varphi_A$ | $0.83 \cdot \left(\frac{r}{R}\right)^{5.21} + 0.17$ |
| Mass transfer coefficient during incubation, $h_{m,inc}$ | $4 \times 10^{-10} \text{ m} \cdot \text{s}^{-1}$ |
| Mass transfer coefficient during clearance, $h_{m,cl}$ | $8 \times 10^{-7} \text{ m} \cdot \text{s}^{-1}$ |
| Clearance time, $t^{\frac{1}{2}}$ | 22 hrs |
| Only for Trastuzumab |  |
| Concentration of total available receptors, $R_T$ | 530 nM |
| Dissociation rate constant, $k_d$ | $4 \times 10^{-3} \text{ s}^{-1}$ |
| $K_D \equiv \frac{k_d}{k_a}$ | 5 nM |
| Internalization rate constant, $k_{int}$ | $1.5 \times 10^{-5} \text{ s}^{-1}$ |

### Modeling the killing efficacy of the radioisotope cargo

The isotope cargo is an α-particle-emitting radionuclide, actinium-225 (^225^Ac), with high cytotoxic efficacy and minimal irradiation of the tissue(s) surrounding the solid tumor(s). Each ^225^Ac emits four α-particles and produces three radioactive daughter isotopes during its decay chain ^13,14^. We assume that cancer cell death follows first-order kinetics:

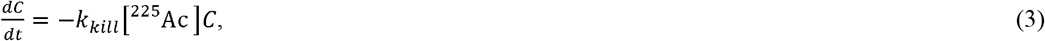

where *c*(*r,t*)is the local density of surviving cancer cells at distance *r* from the center of the spheroid and time, *t*; [^225^Ac] denotes the local concentration of the isotope; *k*_*kill*_ is the cancer cell killing rate constant.

The killing dynamics can be related to the isotope decay rate, 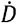. Given that the specific activity of ^225^Ac is ~2000 MBq/nmol, the decay rate is estimated as: 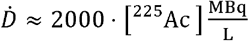, with [^225^Ac] expressed in nM. Integrating equation (3), yields the survival fraction (*SF*) of cancer cells:

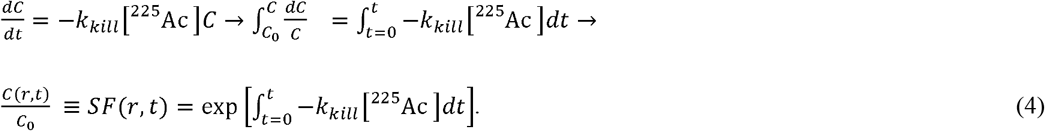

The concentration of ^225^Ac is determined by the spatial distribution of the antibody carrier (Trastuzumab or Rituximab, as model non-binding/low-affinity antibody), in all forms: interstitial, receptor-bound, or internalized. Assuming a constant molar isotope to antibody ratio, *a*_*r*_, the ^225^Ac concentration is given by:

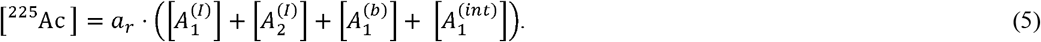

Here, *a*_*r*_ is determined from the specific activity of the conjugate. For Trastuzumab-radioconjugate with a specific activity of 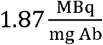 and molecular weight 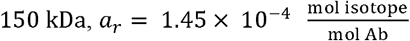. For carrier cocktails, the total ^225^Ac concentration is computed by summing the contributions from all carriers in all forms.

## Data and Code Availability

The data and code supporting the findings of this study will be available upon publication or upon reasonable request.

## Acknowledgements

This study was partly supported by the Congressionally Directed Medical Research Programs, Award number HT9425-24-1-1003 to SS.

